# Stress response of *Chironomus riparius* to changes in water temperature and oxygen concentration in a lowland stream

**DOI:** 10.1101/266528

**Authors:** Alessandro Manfrin, Stefano Larsen, Massimiliano Scalici, Sven Wuertz, Michael T. Monaghan

## Abstract

The increasing impairment of lotic ecosystems has promoted a growing effort into assessing their ecological status by means of biological indicators. While community-based approaches have proven valuable to assess ecosystem integrity, they mostly reflect long-term changes and might not be suitable for tracking and monitoring short-term events. Responses to rapid changes in environmental conditions have been rarely studied under natural conditions. Biomarkers offer the benefit of integrating biological responses at different time scales. Here we used a field experiment to test how the synthesis of heat shock protein 70 (HSP70) and Haemoglobin (Hb) in laboratory-reared larvae of *Chironomus riparius* (Diptera, Chironomidae) were influenced by short-term changes to water temperature and oxygen concentration in a lowland stream. Our aim was to determine whether HSP70 mRNA expression and Hb content could be used as an *in situ* “early warning system” for freshwater habitats undergoing environmental change. HSP70 exhibited a clear response to changes in temperature measured over a one-day period, confirming its suitability as an indicator of environmental stress. Hb concentration was related to oxygen concentration, but not to temperature. Our findings support the hypothesis that depletion in oxygen induces Hb synthesis in *C. riparius* larvae. Because tolerance to low oxygen is not only related to total Hb, but also to a more efficient uptake (binding to Hb, e.g. Bohr effect) and release of oxygen to the cell (Root effect), we cannot discern from our data whether increased efficiency played a role. We suggest that *C. riparius* is a suitable model organism for monitoring sub-lethal stress in the field and that the approach could be applied to other species as more genomic data are available for non-model organisms.

## Introduction

Streams and rivers are among the most threatened ecosystems, having been modified globally by catchment land-use changes, water abstraction, channelization, pollution and invasion of alien species (Vörösmarty et al., 2010; Dudgeon et al., 2006). Additionally, climate change is expected to alter hydrology and temperature regimes with severe effects on organisms and ecosystem functions (Ormerod and Durance 2012; Li et al., 2012; Floury et al., 2013). This increasing impairment of lotic ecosystems has promoted a growing effort into assessing their ecological status by means of biological indicators and sentinel species (Friberg, 2014). The classification of the ecological status of rivers is officially based on the assemblage structure of key taxonomic groups (e.g., Hering et al., 2003; Traversetti et al., 2015). While assemblage-based approaches have been proven valuable in the assessment of ecosystem integrity (Bae et al., 2014), they mostly reflect long-term changes, associated with the local extirpation of sensitive taxa and overall changes in community composition. This approach may not be suitable for identifying and monitoring the effects of short-term events such as droughts and floods or other sub-lethal episodic events, whose frequency and magnitude is expected to increase in the near future (Ledger and Milner 2015). Biomarker assays (i.e. non-lethal responses of biological systems) are often used in eco-toxicological studies to assess the effects of pollutants, but their potential for tracking environmental change in the field has received little attention (Traversetti et al., 2017). Ideally, integrating indicators in a hierarchical fashion, from sub-organismal to organismal, population and community levels (Sures et al., 2015) should improve the assessment of ecosystem health over multiple spatio-temporal scales (Cajaraville et al., 2000; Lagadic et al., 2000; Colin et al., 2016).

A promising approach is to use multiple indicators of stress in organisms (Frank et al., 2013). Multiple biomarkers may produce the benefit of integrating biological responses at different time scales and levels of organisation (Den Besten, 1998; Lagadic et al., 2000; Scalici et al., 2015). Two potential biomarkers for measuring sub-lethal effects in stream macroinvertebrates are heat shock proteins (HSP) and haemoglobin (Hb). HSP70 is a set of chaperon proteins involved in ensuring the correct folding and unfolding of proteins, and its expression is rapidly regulated by changes in physical (i.e. temperature) and chemical conditions (Lencioni et al., 2009; Lee et al., 2006), but it is not affected by handling stress (Sanders, 1993). The expression of HSP70 is therefore considered a short-term “early warning” indicator of environmental changes (Yoshimi et al., 2009; Folgar et al., 2015). For example, Lencioni et al (2013) observed an increase in HSP70 expression after 1h of heat stress at 26 °C in a cold-adapted non-biting midge (Diptera, Chironomidae) larvae.

Chironomidae larvae can be abundant in degraded freshwater habitats, and are thus considered indicators of poor water quality and early colonizers after large-scale disturbances (Serra et al. 2017). Resistance and resilience of chironomids is often attributed to the presence of hemoglobin (Hb), which allows them to tolerate low oxygen concentrations (Moller Pillot, 2009). In *Chironomus riparius,* Choi et al. (2001) observed a 151% increase in total Hb after 24 hours of hypoxia. Chironomidae larvae have been reported to secrete up to 16 different Hb types (Choi and Ha, 2009; Green et al., 1998). Such diversity of Hbs with specific binding properties allows for a fine-tuned loading and unloading of O2 that regulates its delivery to specific tissues under variable environmental conditions (Choi and Ha, 2009; Ha and Choi, 2008; Weber and Vinogradov, 2001). We used a field experiment to test how HSP70 expression and Hb production were influenced by short-term changes to temperature and oxygen concentration in a lowland stream. Laboratory-reared larvae of *Chironomus riparius* (Diptera, Chironomidae), a widespread species considered a model organism in aquatic toxicology (Lee et al., 2006; Lencioni et al., 2009; Morales et al., 2011; Marinkovic et al., 2011), were placed in a stream and sampled over a period of 1 to 8 days, while experiencing rapid variation in temperature and oxygen concentration.

## Methods

The study was carried out in the lowland stream Groote Molenbeek in Limburg, Netherlands. In June 2010, two experimental reaches (upstream, downstream) were designated along the stream, each *ca.* 50 m in length and separated by *ca.* 200 m. In July 2010 the upstream and downstream reaches were separated by an artificial dam and a by-pass was constructed (Fig. 1a, b). The aim was to simulate summer drought conditions in the downstream reach, e.g., reduced water flow, reduced oxygen concentration, and increased water temperature. Experiments were performed in the upstream reach in June, prior to dam construction, and in both upstream and downstream reaches in August, after dam construction. No experiment was conducted in the downstream reach in June because abiotic conditions were nearly identical to those in the upstream reach. In August, heavy rainfall caused large and rapid variations in oxygen and water temperature in both reaches. While this event disrupted the desired effect of the experimental drought, it provided the opportunity to quantify short-term responses to sub-lethal environmental change in all reaches. Therefore, we did not compare control and experimental reaches, but rather we measured the physiological responses of *C. riparius* to the environmental changes experienced *in situ*. The following environmental variables were measured each day at 08:00 throughout the sampling periods in June and August: Temperature (°C), dissolved oxygen (mg O_2_ l^-1^), conductivity (μS cm^-1^), and pH (measured with a Multi 340i/SET immersion probe WTW, Weilheim, Germany), water depth (m) and flow velocity (m sec^-1^) (measured using a 2030 flow-meter; General Oceanics, Miami, USA).

**Figure 1.**
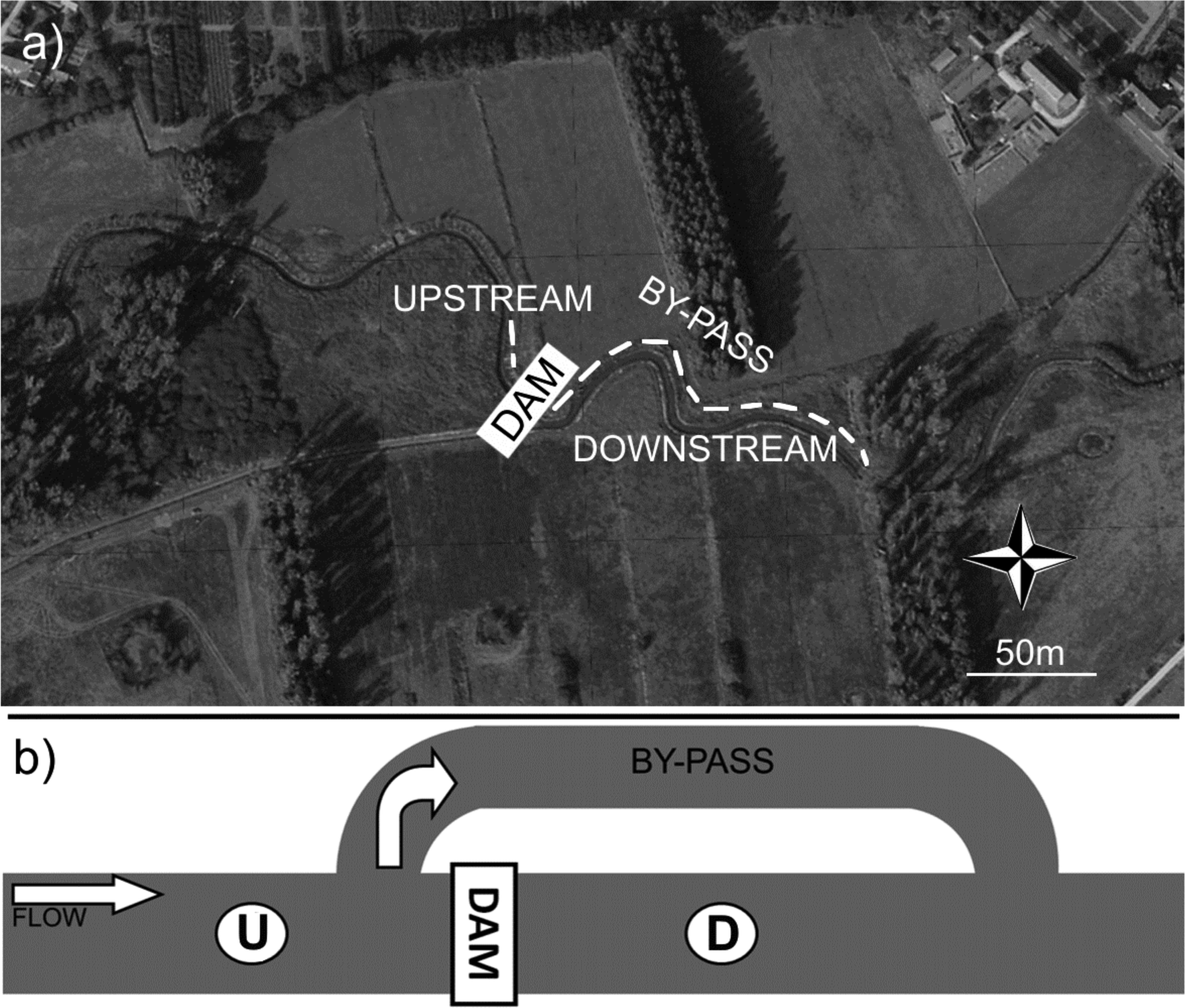
Study site. (a) overview of experimental reaches and dam position on the River Grote Molenbeek (51°23’30.79” N 6°2’31.89” E) (Sevenum, NL); (b) schematic view of dam, by-pass, upstream (U) and downstream (D) sites.

*Chironomus riparius* individuals were obtained from a permanent laboratory population at the Department of Aquatic Ecotoxicology in Frankfurt am Main, Germany. The single origin presumably minimized the genetic diversity among individuals (Nowak et al., 2012). Eggs were shipped to the IGB in Berlin, and after hatching, larvae were reared in aquaria for 4 months prior to the experiment according to the OECD (2004) guidelines. Laboratory aquaria were filled with fine quartz sand as substrate. Aquaria were constantly aerated and kept in a climate chamber in controlled conditions (20 °C, light:dark 16h:8h). Larvae were fed with commercial TetraMin^®^ fish food (Tetrawerke, Melle, Germany). Mesh cages (16 × 12 × 12 cm; mesh: 0.2 mm; Fig. 2a) were designed ad-hoc from aquarium isolation chambers (Hagen Marina, Montreal, Canada) and used to transfer *C. riparius* larvae from the laboratory to the field and to introduce larvae into the experimental reaches. This procedure enabled rapid sample collection, thus minimising handling stress.

At the start of experiments, 25 mesh cages, with 100 larvae each (third and fourth instar), were placed on the stream bottom (Fig. 2b) in each reach (upstream in June, upstream and downstream in August). Fourth-instar larvae were used for HSP70 expression analysis (collected after 24, 96 and 192 hours of exposure) and Hb analyses (24, 48, 96 and 192 hours of exposure; sample sizes in Appendix 1). Larvae were removed from cages with forceps, placed in cryo vials (Eppendorf), immediately placed in liquid nitrogen and stored at −80°C until analysis.

**Figure 2.**
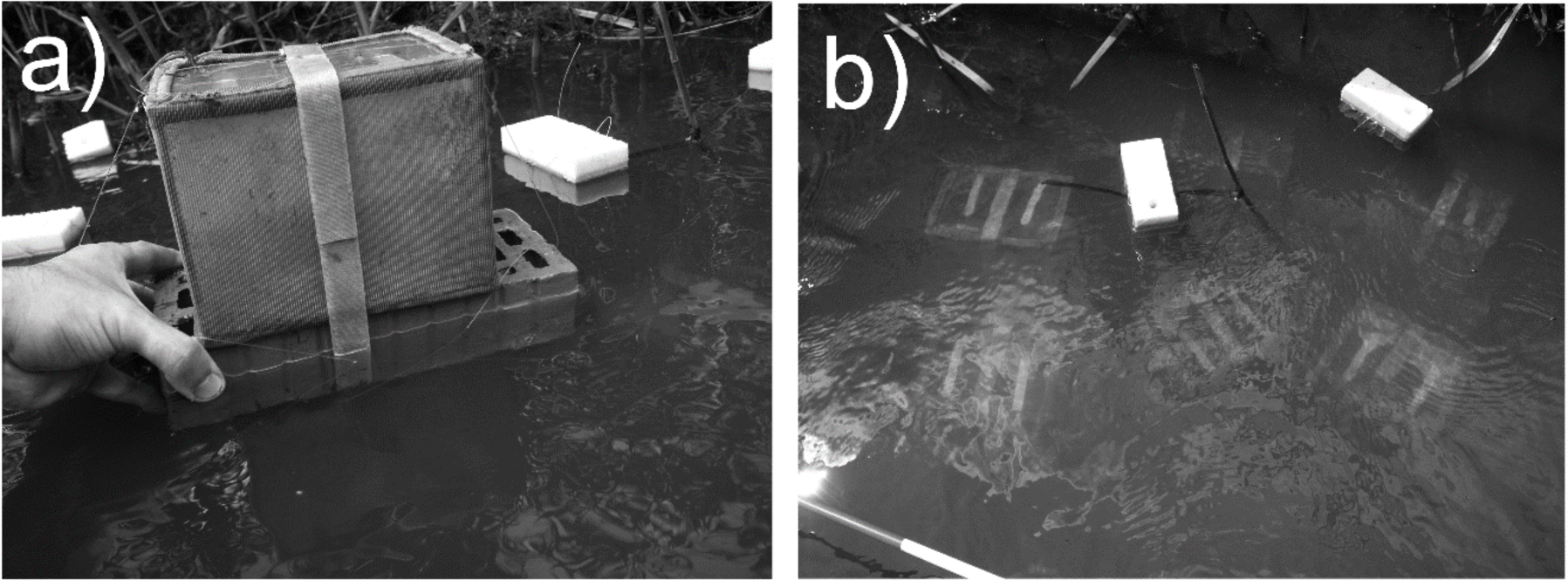
Experimental steup in the stream: (a) *Chironomus riparius* experimental cages (16 × 12 × 12 cm), and (b) their positioning on the river bed in a 100 × 250 cm are in upstream and downstream sites (see Fig. 1).

Total RNA was extracted from 7 individuals per time point and reach (n = 63; Appendix 1) using a Rneasy Mini kit (Qiagen, Hilden, Germany) with on-column DNase digestion (Trubiroha et al. 2009). RNA concentration was measured using a Nanodrop ND-1000 (Thermo Fisher Scientific, Darmstadt, Germany). Reverse transcription was carried out with Affinity Script transcriptase (Agilent/Stratagene, Waldbronn, Germany). Primers for HSP70 and ß-actin (Appendix 2) were designed using data from Park et al. (2010) and Morales et al. (2011) and specificity was confirmed by direct sequencing. Quantitative PCR was carried out with a Mx3005 (Agilent/Stratagene) using hot start polymerase (Phire Taq II, Life Technologies) and SYBR Green in a 20 µL reaction volume (2 µL diluted cDNA, 375 nM of each primer, 1x Taq buffer, 2 mM MgCl2, 0.5 mM each dNTP, 0.5 fold diluted SYBR-Green I solution, 1 U polymerase) under the following conditions: 98°C initial denaturation for 4 min, followed by 40 cycles of 98°C denaturation for 20 s, 62°C primer annealing for 15 s, and 72°C extension for 20 s. PCR efficiencies were determined in triplicate with a dilution series of pooled cDNA (ß actin 99.6%; HSP70 98.4%). All samples were determined in duplicate. Expression was determined by the comparative ΔΔC _T_method (Pfaffl, 2001) with ß actin used as a baseline (housekeeping) gene considering a calibrator sample (pooled cDNA) and correction for efficiency. Specificity of amplification was monitored by melting curve analysis.

Total Hb was measured in nine individuals per time point and reach (n=108; Appendix 1) using the cyanomethemoglobin method with a diagnostic haemoglobin reagent (DiaSys, International, Holzheim, Germany) as described by Wuertz et al. (2013). All samples were measured twice with an Infinite 200 microplate reader (Tecan, Mainz-Kastel, Germany) at 540 nm and concentration was calculated using a standard dilution series (120 mg/L haemoglobin standard, Diaglobal GmbH, Berlin, Germany). Total Hb was normalized to the total protein concentration determined by the Bradford (1976) method (RotiQuant Kit, Germany) as µg Hb/µg total proteins.

Linear mixed-effect (LME) models were used to analyse variation in HSP70 expression and Hb concentration in relation to variation in environmental conditions.

After testing for collinearity using a Spearman test (also from among all the measured environmental variables, see Appendix 3) the model for both HSP70 and Hb initially incorporated T, O _2_and changes in temperature (ΔT) and oxygen (ΔO_2_) as fixed factors. The latter two variables were calculated as the absolute change (i.e., increase or decrease) in T (°C) or O _2_(mg O_2_ L^-1^) from the previous sampling time (every 24h). Model factors were then backward-selected using likelihood ratio tests against reduced models (without the fixed factor) (Zuur et al. 2009). Finals models included fixed factors ΔT, ΔO _2_ and T for HSP70 model and O_2_ for Hb model. Collection time nested in the reach-season (upstream-June; upstream-August; downstream-August) was considered as a random factor to account for repeated sampling. The variance explained by each model was calculated as marginal (R^2^m) (Nakagawa and Schielzeth, 2013) using the MuMln package (Barton 2016) for R vn 3.3.1 (R Core Team, 2015). Residuals were tested for normality with a Wilk-Shapiro test and qq-plots, and scores were log-transformed to remove heteroscedasticity if necessary.

## Results

Initial environmental conditions in June (Fig. 3) were assumed to be identical between the upstream and downstream sites in August due to their close proximity and the lack of a dam. In August, mean water flow, channel depth, dissolved oxygen, and conductivity varied after placement of the dam and after heavy rain events (Appendix 4b, c, e, f). Flow, depth, and conductivity all decreased in August and varied between reaches (Appendix 4c, f), whereas oxygen increased substantially in the upstream reach and less in the downstream reach, compared to June (Fig. 3a, Appendix 4b). HSP70 expression in the upstream reach in June was stable after 24 and 96 hours but increased at 192 hours in June. Expression peaked at 96 hours in both reaches in August, following a rapid change in T (Fig. 3b). The mixed effect model combining all data from June and August indicated significant positive relationships between HSP70 expression and change in temperature (ΔT) and oxygen (ΔO_2_) over the previous 24 hr (Table 1).

**Table 1.**
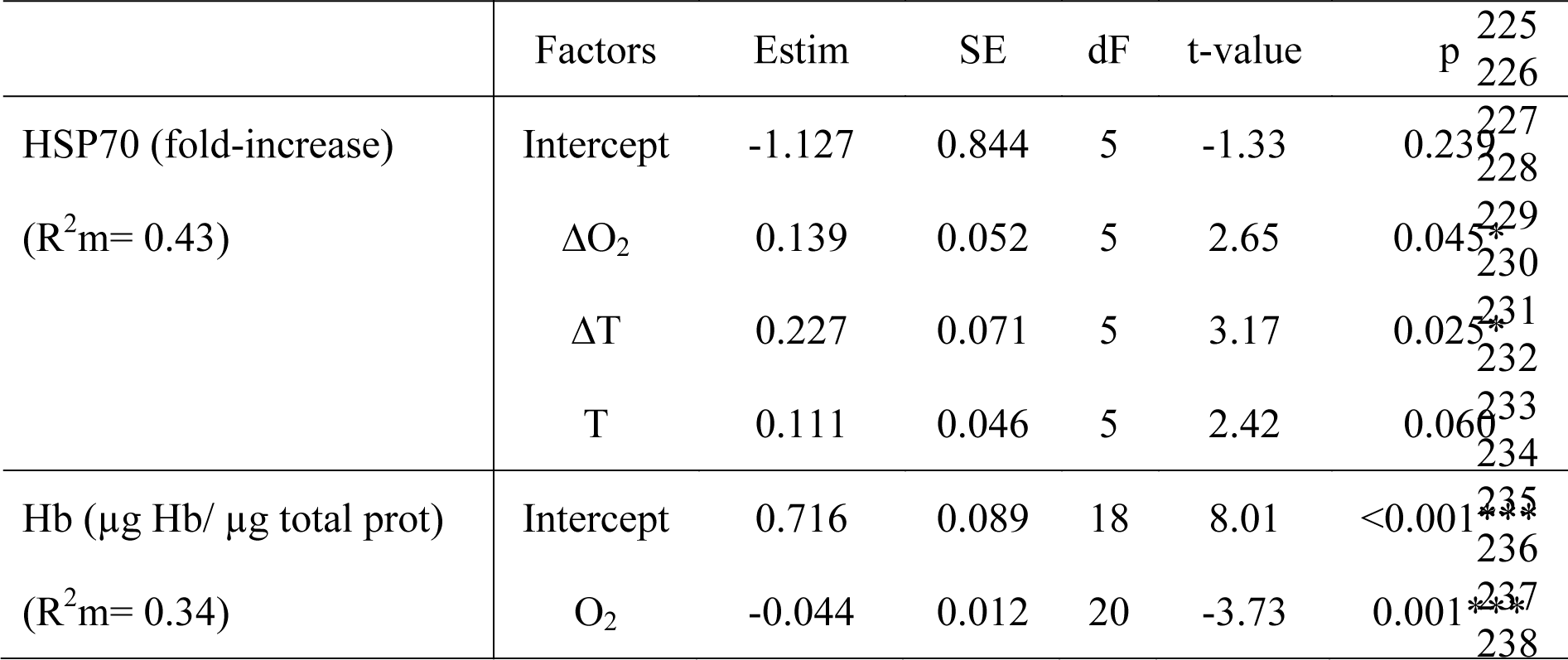
Results of linear mixed effect (LME) models for the relative heat shock protein 70 (HSP70) expression (standardized to the calibrator) and hemoglobin (Hb) response to environmental changes, including marginal variance (R^2^m), estimate of the fixed effects (Estim), standard error (SE), degrees of freedom (dF) and t-statistic (t-value and factor significance). Fixed factors: change in Oxygen (ΔO_2_, mg O_2_ l^-1^); absolute change in T (ΔT, °C); Temperature (T, °C); oxygen (O_2_, mg O_2_ l^-1^). * = *p*< 0.05, ** = *p*< 0.01, *** = *p*< 0.001.

**Figure 3.**
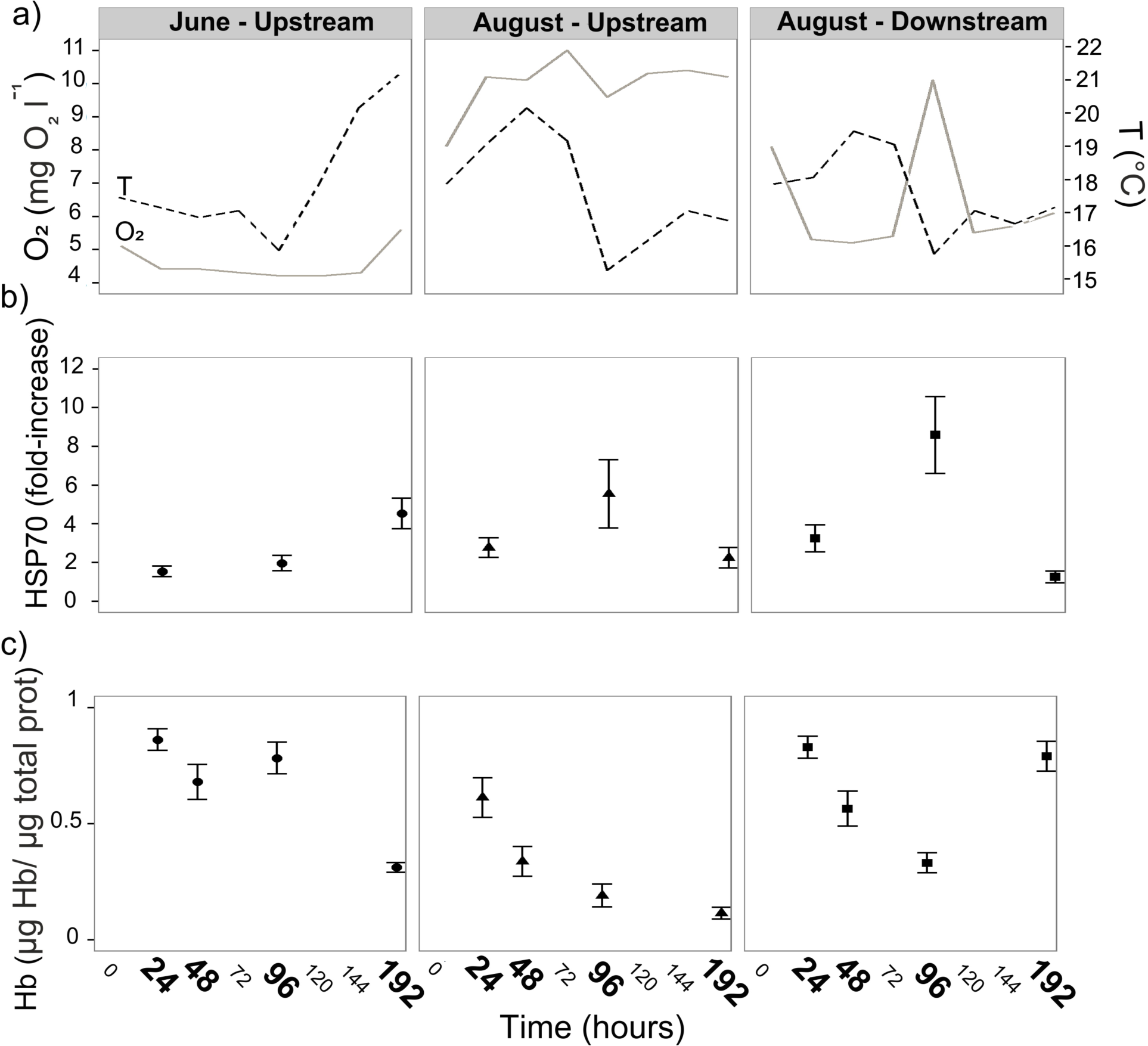
Environmental variables and physiological responses of *Chironomus riparius* larvae: (a) changes in oxygen concentration (solid line) and temperature (broken line) as measured at 08:00 each day during the experiments in June and August and at 1 hour before sample collection (hours indicated in bold on the y-axis); (b) mean (± SE) relative heat shock protein 70 (HSP70) expression standardized to the calibrator at 24, 96, and 192 hours; (c) mean (± SE) haemoglobin (Hb) concentration at 24, 48, 96 and 192 hours.

Hb concentrations in the upstream reach remained similar after 24, 48, and 96 hours and were lowest after 192 hours in June, after a mild increase in dissolved O2. (Fig. 3c). In August, Hb declined steadily over 24, 28, and 96 hours in both reaches. In the upstream reach, Hb continued to decline after 192 hours during relatively stable O2 levels, but increased markedly in the downstream reach following a peak and subsequent rapid decline in O2 after in 192 hours (Fig. 3c). The mixed effect model indicated that Hb content increased with decreasing oxygen concentration (O_2_) (Table 1).

## Discussion

We applied an eco-toxicological stress-response approach to a field experiment in order to examine how changes in water temperature and dissolved oxygen concentration influenced two physiological biomarkers in a model organism. Our markers were chosen to reflect short-term (HSP70 mRNA expression) and medium-term (blood Hb content) responses to environmental changes. Contrary to conventional bio-assessment programmes, where the presence or abundance of different aquatic organisms are used as indicator of environmental change or degradation, our aim was to determine whether pysiological biomarkers in a model organism could be used *in situ* as an “early warning system” for freshwater habitats undergoing environmental change. Assemblagelevel responses may manifest only at a later stage of environmental degradation, thus hindering prompt mitigation actions. Many studies of stress response are conducted in the laboratory and often under conditions unlikely to represent those of natural habitats (see Sures et al. 2015). Our intent was to extend this approach to realistic field conditions. In addition, we used a model organism group (Chironomidae) that is almost ubiquitous in aquatic habitats and is among the first colonizers after disturbance events such as droughts or floods (Calle-Martinez and Casas, 2006; Langton and Casas, 1998; Marziali et al., 2010; Punti et al., 2007). HSP70 exhibited a clear response to changes in temperature (and partially in oxygen) measured over a one-day period prior to sampling. The expression of HSP70 is known to be related to acute cellular stresses (Morimoto and Santoro 1998), and Feder and Hofmann (1999) observed the presence of HSP-inducing microhabitats (e.g. shallow or stagnant water systems) where mild environmental variations (e.g. temperature) induced variations in HSP expression. HSP70s have also been reported reliable means of detecting such stress (Lencioni et al., 2009, Foster et al., 2015). This may be due to the fact that in dynamic systems such as small waterbodies, environmental parameters like temperature and oxygen vary slightly but continuously, causing an increase of the long-term memory formation as an adaptive response of the organisms (Foster et al., 2015). Memory formation increases synaptic efficacy and improves the adaptive responses to stress conditions including the basal mRNA transcriptional system (Stork and Welzl, 1999, Monari et al., 2011) in which HSP70 are also included. This further supports the suitability of the HSP70 as multi-stressors indicator.

In this study, haemoglobin concentration was related to oxygen concentration, but not to water temperature. Results from other studies indicate that the tolerance of *C. riparius* larvae to low levels of dissolved oxygen is associated with increased heamoglobin in their hemolymph (Weber, 1980; Choi et al., 2000), which allows sustaining aerobic and anaerobic metabolism (alcoholic fermentation) at the same time for short periods (Frank, 1983). Under hypoxic conditions, Hb synthesis is stimulated and used for aerobic metabolism (Choi et al., 2000; Lee et al., 2006; Rossaro et al., 2007). This process likely occurred in our experiment where the observed depletion in oxygen concentration induced synthesis of Hb in *C. riparius* larvae. Nonetheless, tolerance to low oxygen is not only related to total Hb, but also to a more efficient uptake (binding to Hb; Bohr effect) and release of oxygen to the cell (Root effect). However, we cannot discern from our data whether increased efficiency played a role. The synthesis of HSP70 and Hb are likely linked, because temperature and oxygen concentration are closely interconnected. Our results suggest that the responses of HSP70 and Hb to environmental change represent an integrated process in which HSP70 increased as a direct consequence of increased temperature. Subsequently, increased temperatures likely led to a decline in Oxygen concentration that promoted additional synthesis of Hb.

In conclusion, we suggest that the sub-lethal stress response at multiple markers make *C. riparius* a suitable biological tool for the assessment of short-term, sub-lethal effects of environmental change in the field. The different temporal scales involved in the response of the two markers indicate that a variety of impacts could be assessed prior to local extinction. Because the frequency of extreme hydrological events is likely to increase in the future owing to global climate change, ‘early-warning’ indicators could allow the rapid assessment of environmental degradation. As more genomic data are made available, our approach could be extended to other taxonomic groups with different environmental requirements and additional genetic markers.

## Acknowledgements

We thank Annemarie Garssen (University of Utrecht) for help in the field, Marino Marinkovic (University of Amsterdam) and Karsten Nowak (Senckenberg Research Institute) for help with *Chironomous* larvae, and Wibke Kleiner, Katrin Preuss, and Ann-Christin Honnen (IGB) for help in the laboratory. Furthermore, we thank Bernd Sures (University of Duisburg-Essen) for valuable comments on a previous version of the manuscript. AM was partially funded by the Education, Audiovisual and Culture Executive Agency of the European Commission within the Erasmus Mundus Joint Doctorate Program SMART.

## Supplementary material

**Appendix 1.** Sample sizes (number of individual *C. riparius*) for each analysis in the study given for each time point of collection in hours (h), with HSP70 = heat shock protein 70 mRNA expression; Hb = haemoglobin concentration.

| Reach | Marker | Number of samples (*n*) |  |  |  |  |
| --- | --- | --- | --- | --- | --- | --- |
|  |  | 24h | 48h | 96h | 192h | total |
| June – upstream | HSP70 | 7 |  | 7 | 7 | 21 |
|  | Hb | 9 | 9 | 9 | 9 | 36 |
| August – upstream | HSP70 | 7 |  | 7 | 7 | 21 |
|  | Hb | 9 | 9 | 9 | 9 | 36 |
| August – downstream | HSP70 | 7 |  | 7 | 7 | 21 |
|  | Hb | 9 | 9 | 9 | 9 | 36 |

**Appendix 2.**
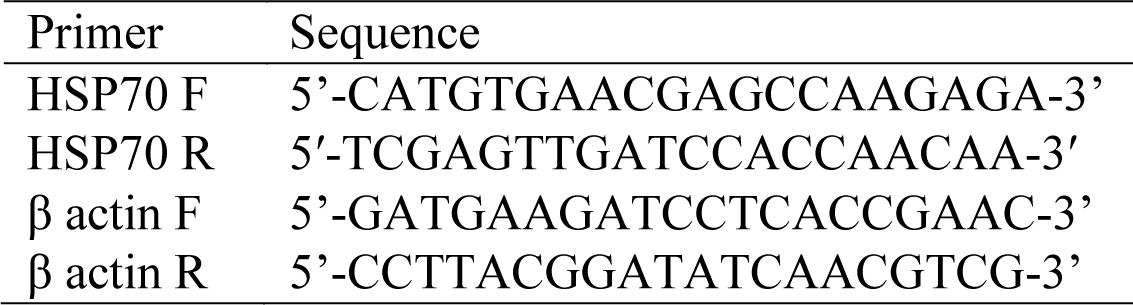
Newly designed forward (F) and reverse (R) primer sequences used for the RT- PCR for heat shock protein 70 (HSP70) and β actin gene expression in *C. riparius*.

**Appendix 3.**
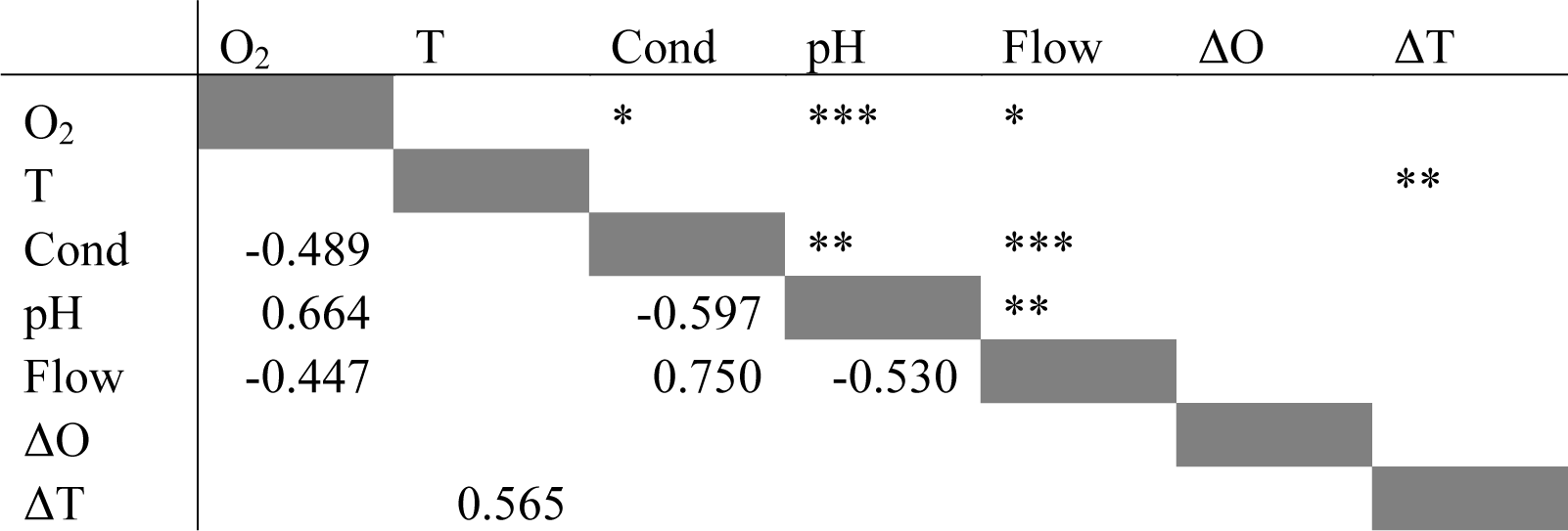
Statistics from Spearman correlation test (scores under diagonal) and p-values (over diagonal) for correlated environmental variables: oxygen (O_2_, mg O_2_ l^-1^); temperature (T, °C); conductivity (Cond, μS cm^-1^); pH; water flow (Flow, m s^-1^); change in Oxygen (ΔO_2_, mg O_2_ l^-1^); changes in T (ΔT, °C).

**Appendix 4.**
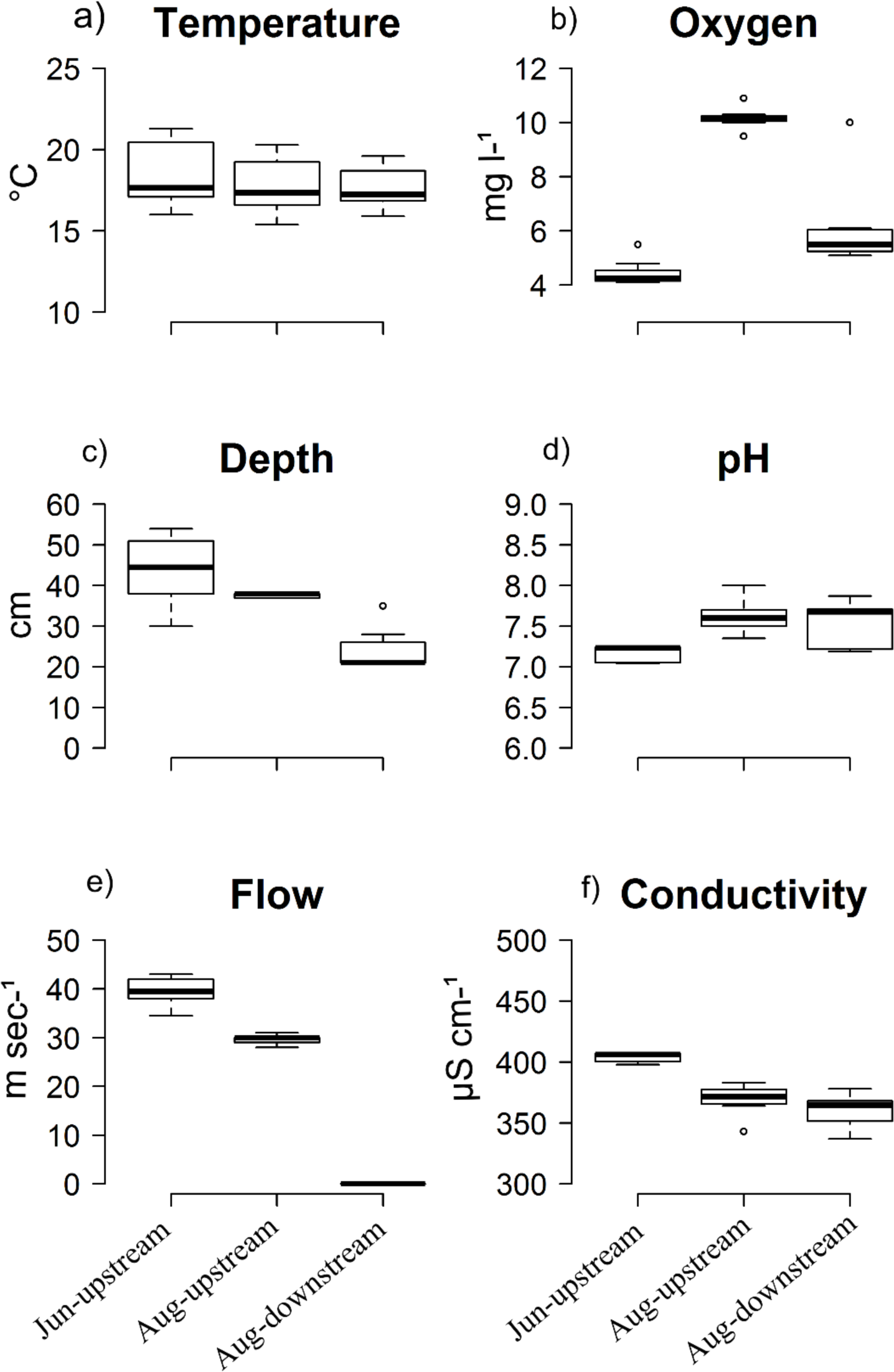
Environmental variables (panels a to f) (median, first and third quartile, minimum and maximum) from data collected every 24 hours at 08:00 from 24h to 192 h of the experiments) in the two reaches in June and August.

